# Unraveling molecular interactions in a phase-separating protein by atomistic simulations

**DOI:** 10.1101/2020.05.16.099051

**Authors:** Matteo Paloni, Rémy Bailly, Luca Ciandrini, Alessandro Barducci

**Affiliations:** Centre de Biochimie Structurale (CBS), INSERM, CNRS, Université de Montpellier, Montpellier, France

## Abstract

Membraneless organelles are dynamical cellular condensates formed by the liquid-liquid phase separation of proteins and RNA molecules. Multiple evidence suggests that disordered proteins are structural scaffolds that drive the condensation by forming a dynamic network of inter- and intra-molecular contacts. Despite the blooming research activity in this field, the structural characterization of these condensates is very limited and we still do not understand how the phase behaviour is encoded in the amino-acid sequences of the scaffolding proteins. Here we exploited explicit-solvent atomistic simulations to disentangle the molecular interactions governing the phase behaviour of the N-terminal disordered region of DEAD-box helicase 4 (NDDX4), which is a well-established model for phase separation *in vitro* and *in vivo*. Single-molecule simulations clarified the interplay between the intramolecular interactions that shape NDDX4 conformational ensemble and the known determinants of its phase behaviour, such as the attraction between oppositely-charged regions and the presence of arginine and phenylalanine. We then investigated intermolecular interactions associated with phase separation via a divide-and-conquer strategy based on the simulations of various NDDX4 fragments at high concentration. Our approach allowed us to probe conditions mimicking real condensates and revealed, in agreement with mutagenesis results, how these interactions arise from the complex interplay of diverse molecular mechanisms. Particularly, we characterized the transient formation of clusters of arginine and aromatic residues, which may stabilize the assembly of several MLOs. Overall, our results reveal the potential of atomistic simulations in the investigation of biomolecular phase separation paving the way for future studies.

## Introduction

While cellular compartmentalization is canonically thought to rely on membrane-bound organelles, such as the nucleus or mitochondria, biomolecular condensates devoid of lipid membranes are currently recognized to orchestrate a variety of biological processes and have attracted growing interest in the last decade^1–4^. These membrane-less organelles (MLOs) are dynamical supramolecular structures with liquid-like properties that emerge from the condensation of a variety of biomolecular components. An increasing amount of evidence suggests that MLOs are formed via liquid-liquid phase separation (LLPS) processes driven by multivalent protein-protein and protein-nucleic acid interactions^5–7^. As MLOs are currently thought to play a major role in organizing the cellular environment, unveiling the still elusive physico-chemical principles that underlie their assembly and regulation has paramount importance to improve our understanding of the molecular basis of human health and disease^8^.

In this challenge, a key breakthrough was achieved by identifying non-dispensable protein components of various MLOs, such as LAF-1^9^, DDX4^10^, FUS^11^, hnRNPA1^12^, and TDP-43^13^, which self-assemble *in vitro* into condensed droplets reminiscent of the corresponding cellular structures. These proteins are assumed to act as molecular scaffolds that drive LLPS in the cellular environment and they hence represent useful minimal models to investigate the properties of MLOs through controllable *in vitro* experiments. Remarkably, all these scaffold proteins share some analogies, such as a remarkable degree of structural heterogeneity due to the presence of low-complexity, disordered domains, which are necessary for *in vitro* and *in vivo* phase separation^9–11^. Furthermore, pioneering NMR studies indicate that this conformational flexibility is preserved even upon LLPS and that the condensed phase is thus stabilized by a dynamical network of transient inter-chain interactions^14,15^.

These findings stimulated several investigations aiming to elucidate the relationship between the amino-acid sequences of these intrinsically disordered proteins/regions (IDPs/IDRs), their conformational properties at the single-molecule level, and the propensity to undergo liquid-liquid demixing. While we are still far from a complete understanding of how the phase behavior is actually encoded in the protein sequence, recent experimental and computational results have shed some light on the molecular grammar governing cellular LLPS^16,17^. Particularly, a few studies investigated the long-range electrostatic interactions between charged amino-acids in IDP/IDRs: polymer theory has been used to rationalize how the patterning of charged residues in the sequence affects both the conformational ensemble of isolated proteins and their phase separation properties^18–20^. Furthermore, several evidence indicate that amino-acids with aromatic (Phe, Tyr) or large-sized, unsaturated side-chains (Arg, Gln) are key determinants of *in vivo* and *in vitro* LLPS^10,14,17,21–24^ and short-ranged attractive forces due to π-π or cation-π interactions have been invoked to rationalize these observations^25^.

Molecular simulations are a powerful and promising approach for dissecting the driving forces of LLPS at the molecular level and for accessing the structural details of the condensates, which are elusive to most experimental techniques. Particularly, Coarse-Grained (CG) models with a one-bead-per-residue resolution, based on Debye-Huckel electrostatics and empirical contact potentials, have been shown to qualitatively capture sequence-dependent properties of IDP/IDR^26–29^. Recently, this strategy has been successfully applied for determining coexistence curves of model phase-separating systems^30–32^, for rationalizing the role of phosphorylation on LLPS^33^ and for investigating the correlation between single-molecule conformational ensembles and phase behavior^34^. Furthermore, coarser approaches have been adopted to model the internal organization of heterogeneous MLOs^35–37^. Even if extremely instructive, all these studies invariantly relied on low-resolution, phenomenological CG potentials, which inherently prevented a detailed characterization of the physical interactions driving LLPS. Conversely, the contribution of high-resolution, atomistic simulations to this field has been extremely limited^15,24,38^. This unfortunate shortcoming is mainly due to the demanding computational requirements of this approach, which make the direct simulation of LLPS unfeasible with present-day computers. Furthermore, the accuracy of atomistic force-fields in this context has yet to be carefully assessed, considering the debate on their capability of correctly reproducing the conformational properties of IDRs/IDPs^39–41^ and a recent study that suggested a significant underestimation of stacking interactions^25^.

Here we show that simulations based on last-generation force-field and a ‘divide-and-conquer’ protocol can decipher the intra- and inter-molecular interactions involved in cellular LLPS. We applied this protocol to a model phase-separating system such as the N-terminal disordered region of DDX4 protein (NDDX4), which is responsible for the formation of nuage bodies and has been extensively characterized by structural and biophysical experiments^10,14,42^. Our results provide an accurate, high-resolution picture that is fully consistent with experimental results and pave the way for elucidating at the atomistic level the complex interrelationship between amino acid sequence, single-molecule conformational properties and phase behaviour.

## Results

We first investigated the conformational properties of NDDX4 by simulating a single protein in explicit solvent with a last-generation atomistic force field, which has been validated against experimental data for a set of IDPs^40^. The analysis of 10 MD trajectories in the 1-μs timescale (Fig. 1A) indicated that NDDX4 is a highly flexible molecule that transiently populates a variety of diverse conformational states, in agreement with the picture provided by NMR spectroscopy^10,14^. While an exhaustive characterization of this structural heterogeneity is beyond the current capabilities of explicit-solvent simulations for a large-sized IDP, such as NDDX4 (236 residues), we can profit of 10 μs of aggregated simulation time to safely draw some general conclusions about the protein conformational propensities. Particularly, the simulated conformational ensemble is rather compact (<R_gyr_> = 2.9+-0.3 nm) and it greatly differs from the predictions obtained with statistical IDP/IDRs models based on random-coil approximation (<R_gyr_> ∼ 4.2 nm)^43,44^ (Fig. 1B). Nevertheless, this limited expansion is fully consistent with available experimental data, as proven by the excellent agreement between the protein hydrodynamic radius predicted by MD trajectories (<R_hyd_> = 3.20+-0.16 nm) and the experimental estimate by NMR diffusion measurements (<R_hyd_> ∼ 3.16 nm)^14^. Additionally, the observed compaction of NDDX4 cannot be ascribed to secondary structure elements, whose formation is very limited (Fig. S1), and it rather originates from an extensive network of transient tertiary contacts. In order to characterize these interactions, we first report in Fig.1C a locally-averaged probability *P*_*loc*_*(i)* of forming intra-chain contacts for each residue *i* along the NDDX4 sequence. We notice that while the entire protein is significantly involved in tertiary contacts, a few regions display a particularly high interaction propensity. Interestingly, one of these hotspots, the fragment 132-164, is actually known to play a critical role in liquid-liquid phase separation, as splicing variants of DDX4 lacking this region do not form organelles in physiological conditions *in vitro*^10^. Furthermore, we notice that the intra-chain contact probabilities of arginines in RG/RGG motifs (<P_loc_(i)>=0.81+-0.03), which are a distinctive and conserved feature of many MLO-forming proteins^45^, are significantly higher than the average P_loc_(i) (0.73+-0.02) and of Arg residues not involved in RG/RGG motifs (<P_loc_(i)>=0.76+-0.02).

**Fig.1.**
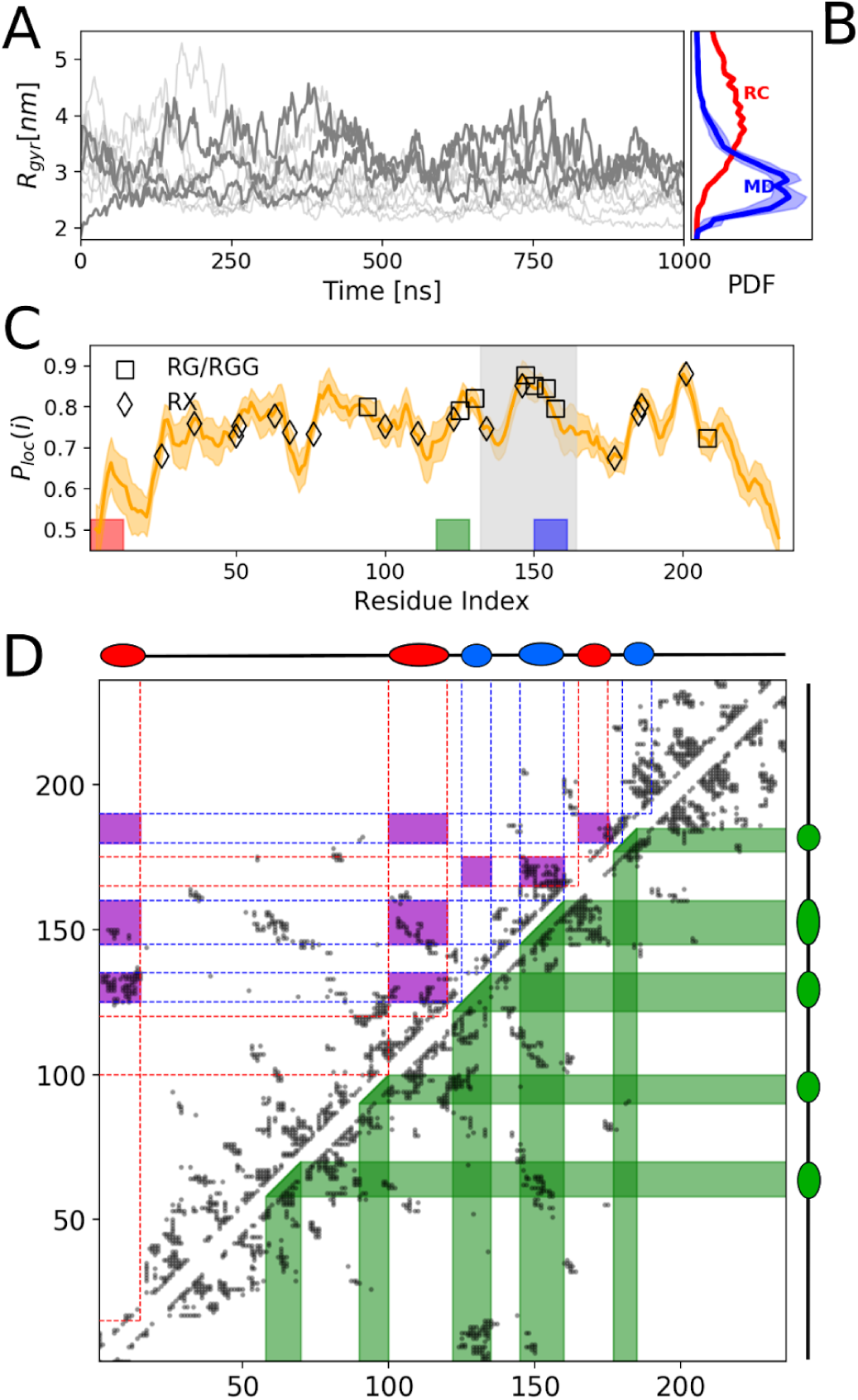
**(A)** Evolution of the radius of gyration (R_gyr_) of NDDX4 in the ten 1-μs MD simulations; for clarity three representative replicas have been highlighted. **(B)** Comparison of the probability density function (PDF) of R_gyr_ from the MD simulations (blu line) with that of a random-coil (RC) model (red line). Shadowed area indicates the error of the mean over ten independent replicas. **(C)** Locally-averaged probability of formation of intramolecular contacts. Error bars represent error of the mean over 10 independent replicas, symbols indicate the position of RG/RGG motifs (squares) and other arginines (diamonds). The grey-shaded area highlights region 132-164, which is necessary for phase separation. Colored squares indicate the position of the 12-residue-long peptides selected to study intermolecular interactions. **(D)** Residue-residue contact probability map. Scatter plot represents the 5% most frequent contacts observed in the MD simulations (see Fig. S2 for the 10% and 15% most frequent contacts). Top left: map of the electrostatic interactions between positively and negatively charged blocks of the protein. Position of the blocks is indicated along the top x-axis, respectively in blue and red. The violet areas highlight the region of favourable electrostatic interactions between oppositely-charged patches. Bottom right: interactions involving segments rich in phenylalanine and arginine. The positions of these fragments are shown along the right y-axis.

Furthermore, mutagenesis experiments^14^ have revealed that LLPS of NDDX4 is hampered by *i)* the weakening of electrostatic interactions due to the scrambling of the patches of acid or basic residues present in the wild-type sequence and by *ii)* the global substitution of phenylalanine with alanine or arginine with lysine. Here we tried to take advantage of these experimental results to investigate the correlation between the driving forces of NDDX4 compaction and the molecular determinants of its phase behaviour. To this aim, we report in Fig. 1D a contact map encompassing the residue-residue interactions most frequently observed in our simulations. A visual inspection of these contacts intuitively indicates that oppositely-charged patches in NDDX4 sequence considerably interact at the single-molecule level, thus favouring spatial proximity of protein regions that are distant in the sequence (top-left half of Fig.1D). Furthermore, fragments of NDDX4 that are particularly rich in phenylalanine and arginine residues exhibit a strong propensity to interact and account for a large fraction of the most favourable intrachain contacts (bottom-right half of Fig.1D). Altogether, our simulations therefore suggest that the molecular determinants of NDDX4 phase separation, i.e. charge-charge attractions and phenylalanines/arginine residues, largely contribute to the organization of the single-molecule conformational ensemble. This observation based on a detailed atomistic description thus corroborates the correspondence between intra- and inter-molecular interactions in phase-separating systems previously suggested by lower-resolution CG simulations^23,32,34^.

Reassured by the agreement of our single-molecule simulations with available data, we then extended our MD strategy to multi-protein simulations to directly probe the intermolecular interactions that drive the phase separation of NDDX4 and stabilize the protein condensates. Unfortunately, the large size and slow dynamics of NDDX4 chains currently prevent a satisfactory description of the protein condensed phase by explicit-solvent atomistic MD. We circumvented this difficulty by focusing on the intermolecular interactions formed by 12-residue-long polypeptides, which correspond to diverse fragments of NDDX4. These fragments, namely Frag1, Frag2, and Frag3, have been selected according to their intra-chain contact propensity and their sequence features (Table 1). Particularly, both Frag1(150-161) and Frag2(117-128) correspond to highly-interacting regions which form promiscuous contacts within the single-molecule NDDX4 ensemble. Frag1 is enriched in RG/RGG motifs and it is characterized by positive net charge, whereas in Frag2 two acidic residues counterbalance the two positively-charged arginines. Conversely, Frag3(1-12) corresponds to the negatively-charged N-term of NDDX4 (net charge −5), which selectively binds positively-charged patches in the sequence but it scarcely interacts with the rest of the protein. We made use of large set of 1-microsecond-long multi-chain trajectories to obtain an exhaustive characterization of homotypic interactions in Frag1, Frag2, Frag3 peptides and heterotypic interactions in Frag1+Frag3 mixtures. In all the cases, 10 peptide chains were simulated in a periodic box representative of a concentrated solution (∼150 mg/ml). In these conditions, the polypeptides form extensive yet highly dynamical intermolecular contacts and transiently explore a wide range of oligomerization states without undergoing irreversible aggregation (see Fig. 2A). We characterized this structural heterogeneity by introducing an variable *N*_*C*_(i) that measures the intermolecular coordination number for the residue *i* (see Methods). This quantity captures the local fluctuations of protein density in the nanometer range and thus it highlights the diverse local environments sampled within the simulations, which range from isolated chains to highly crowded conditions (Fig. 2B). We thus characterize each simulated system by calculating N_C_(i,t) for all the residues and timeframes and evaluating the resulting overall distribution P(N_C_) (Fig. 2C). Interestingly, the comparison of simulation results with the distribution corresponding to the experimental concentration of NDDX4 condensates (∼380 mg/ml^14^) suggests that the transient clusters observed in MD trajectories may reasonably approximate the physico-chemical environment within real droplets. As a consequence, we expect that our simulations allow probing how protein sequence modulates collective intermolecular interactions in conditions that are relevant for phase-separation. In this respect, the bimodal distribution observed for Frag1, which encompasses both a low-density peak and a higher-density region, can be intuitively explained by a competition between electrostatic repulsion (net charge +3) and short-range attractive forces. The low-density peak is significantly less pronounced in the case of Frag2, which is a polyampholyte with a neutral net charge and higher propensity for intermolecular interactions. Conversely, the electrostatic repulsion seems not to be counteracted by compensatory mechanisms in the case of the acidic Frag3 (net charge −5), which strongly prefers solvated states and minimizes higher-density oligomers. This behaviour is drastically reversed in the binary mixture of the oppositely-charged Frag1 and Frag3, which mimics the interaction between charged patches in NDDX4 condensates and strongly favours highly-packed conformations. Overall, this analysis reveals that the three fragments of NDDX4 display markedly different propensities to coalesce at high concentration. Diverse molecular interactions contribute to this distinctive behaviour that is not exhaustively captured by simple parameters, such as net charge and/or mean hydrophobicity, but it recapitulates the intramolecular contact propensity observed in full-length NDDX4 (see Table 1).

**Table 1.** Properties of the fragments selected from the sequence of NDDX4. Mean hydrophobicity is computed according to Kyte-Doolittle hydrophobicity scale^46^, <P_loc_(i)> corresponds to the average value of the fragment from the MD simulations of NDDX4.

| Name | Sequence (position in NDDX4) | Net charge | Mean hydrophobicity | $\langle P_{loc}(i) \rangle$ |
| --- | --- | --- | --- | --- |
| Frag1 | RGSGCRGGFG (150-161) | +3 | 0.43 | 0.81 |
| Frag2 | CEDNPTNRGGS (117-128) | 0 | 0.30 | 0.76 |
| Frag3 | MGDEDEWEAENP (1-12) | -5 | 0.35 | 0.59 |

**Fig.2.**
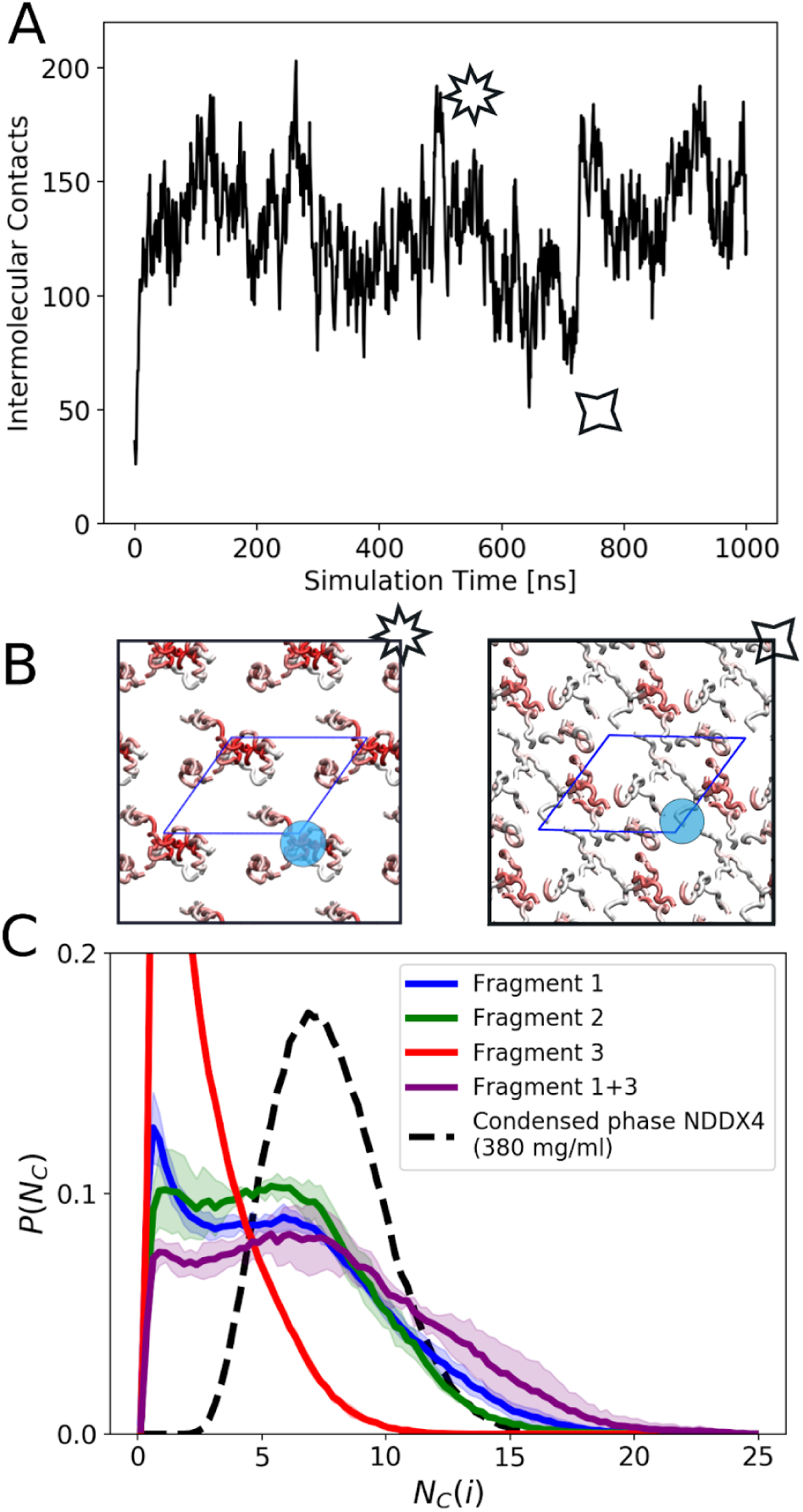
**(A)** Number of intermolecular contacts as a function of simulation time. **(B)** Two representative snapshots from the MD simulations, representing dense (*left*) and solvated (*right*) states. Symbols indicate the corresponding frame and number of intermolecular contacts during the simulation. Protein fragments are drawn in tube representation and each residue is colored by its *N*_*C*_(i) value (high values in red), measuring the number of intermolecular residues within 1 nm, as represented by the blue circle. **(C)** Comparison of the P(N_C_) distributions of the fragments from MD simulations (solid lines) and that of the condensed phase of NDDX4 at ∼380 mg/ml (dashed line). Solid lines represent average distributions, shaded areas indicate standard deviation across the replicas.

Moving beyond this nanoscale characterization, we investigated the intermolecular interactions at higher resolution to better understand the sequence-dependent stabilization of dense phases. Given the close connection between LLPS and amyloid aggregation reported by several studies^11,12,47–49^, we first checked if intermolecular β-structure could account for the high-density clusters observed in our simulations. Conversely, our results indicate that all NDDX4 fragments remained largely disordered at high concentration, with low propensities for secondary structures (Fig. S3). In order to dissect the relevant interactions, we plot in Fig.3 the 2-D maps representing the average probability of intermolecular contact between individual residues in the Frag1, Frag2, and Frag1+Frag3 simulations. Visual inspection of the contact propensities suggests that, despite their limited size, the peptides exhibit rather complex interaction patterns, whose thorough interpretation is extremely challenging. Nevertheless, a few remarkable features can be easily spotted. Particularly, the Frag1 map (Fig. 3A) immediately indicates that the aromatic residues in its sequence (F4 and F11) contribute the most to the intermolecular interactions and they preferentially form Phe-Phe and Phe-Arg pairs. The scenario is more complex in the case of the Frag2 (Fig. 3B), which interacts through distinct mechanisms involving both charged and aromatic residues. Particularly, the arginine-rich RNR motif seems here to represent a molecular pivot for intermolecular contacts as it can strongly interact both with a C-terminal phenylalanine (F11) and with acidic residues in the N-terminal part (E2, D3). In the case of the Frag1+Frag3 mixture, the intermolecular contact probability map (Fig. 3C) is subdivided into three areas accounting for the homotypic interactions between Frag1 (top-right) or Frag3 chains (bottom-left) and heterotypic interactions (bottom-right). As expected, homotypic interactions closely correspond to what observed for the pure Frag1 (Fig. 3A) or Frag3 (Fig. S4) systems but they are significantly less prominent than the contacts between the oppositely-charged Frag1 and Pep3 chains. The latter are stabilized by multiple salt bridges between arginines (Frag1) and glutamate/aspartates (Frag3), although we notice that the most frequent intermolecular contacts (Arg-Trp and Phe-Asn) involve neutral, aromatic residues, even in the case of this electrostatic-driven coacervation. Altogether, the analysis of residue-residue contacts formed by these highly-interacting yet diverse systems indicate that two classes of interactions are prominent in dense multi-chain clusters: the contacts between amino acids with aromatic and/or planar sidechains and transient ionic pairing of oppositely-charged residues. This observation can be confirmed by combining the results obtained for the highly-interacting systems (Frag1, Frag2, and Frag1+Frag3) and noticing that the vast majority of strongest inter-residue contacts falls in one of these two classes (Fig. 3D).

**Fig.3.**
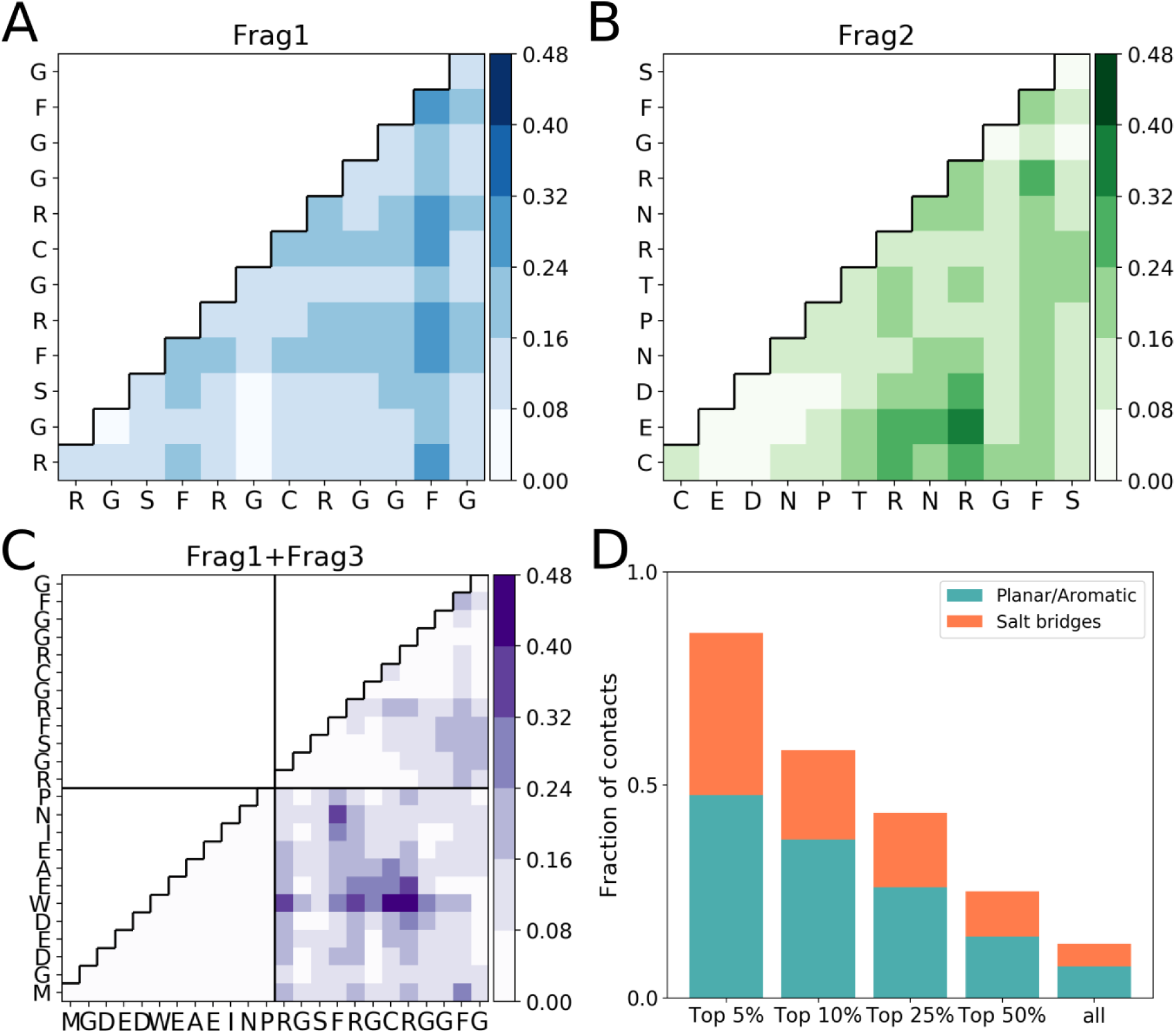
**(A-C)** Average intermolecular contact probabilities of Frag1 (A), Frag2 (B), and Frag1+Frag3 (C) polypeptides. **(D)** Fraction of Planar/Aromatic contacts and Salt bridges for different percentiles of the intermolecular contact probability distribution combining Frag1, Frag2, and the hetero-interaction portion of Frag1+Frag3 simulations. Planar/Aromatic contacts are defined here as Phe-Phe, Phe-Trp, Phe-Asn, Arg-Phe, and Arg-Trp contacts. Salt bridges are defined as contacts between Arg and the negatively-charged Asp and Glu.

In order to further assess the role of specific residues in driving the condensation, we took inspiration from the previously-mentioned mutagenesis experiments^10,14^ and performed additional simulations of the Frag1, Frag2 and Frag1+Frag3 systems where either phenylalanines were replaced with alanines (FtoA mutant), or arginines with lysines (RtoK mutant). We then compared the local density distributions and intermolecular residue-residue contacts for the mutants with those obtained for the wild-type fragments (Fig. 4). In the case of Frag1, intermolecular interactions are weakened both by FtoA and RtoK mutations (Fig. 4A, 4B), even if the former results into a more severe destabilization by abolishing all the critical Phe-Phe and Phe-Arg contacts. Drastic perturbations in local density and interaction patterns are observed also in both Frag2 mutants (Fig. 4C, 4D), although in this system the RtoK substitution is more disruptive because it directly affects the key RNR motif and drastically weakens its interactions with the N-terminal part of the chain. On the contrary, both RtoK and FtoA substitutions have minor impact on the Frag1+Frag3 mixture (Fig. 4E, 4F), as expected for a system that is mostly governed by electrostatic interactions. Altogether, intermolecular interactions between mutant fragments are significantly weaker, or at best equivalent to those observed in wild-type sequences. While far from being exhaustive, our fragment simulations are therefore consistent with the mutagenesis experiments indicating a decreased phase separation propensity of NDDX4 upon FtoA and RtoK substitutions and they can help deciphering the molecular determinants of this behaviour. In this respect, rationalizing the consequences of FtoA replacement is rather straightforward as small-sized alanines cannot substitute for the larger aromatic phenylalanines, which play a key role in the intermolecular interaction network. The interpretation of RtoK substitution is less trivial given the analogies between arginine and lysine, which share the same net charge and similar size. However, our analysis supports the hypothesis that arginines provide a larger contribution to intermolecular contacts by pairing more favourably with aromatic and/or planar residues. Strikingly, this differential behaviour may eventually result also in an effective stabilization of arginine-based salt bridges within the heterogeneous environment observed at high protein concentration, as in the case of Frag2 and Frag1+Frag3 simulations.

**Fig.4.**
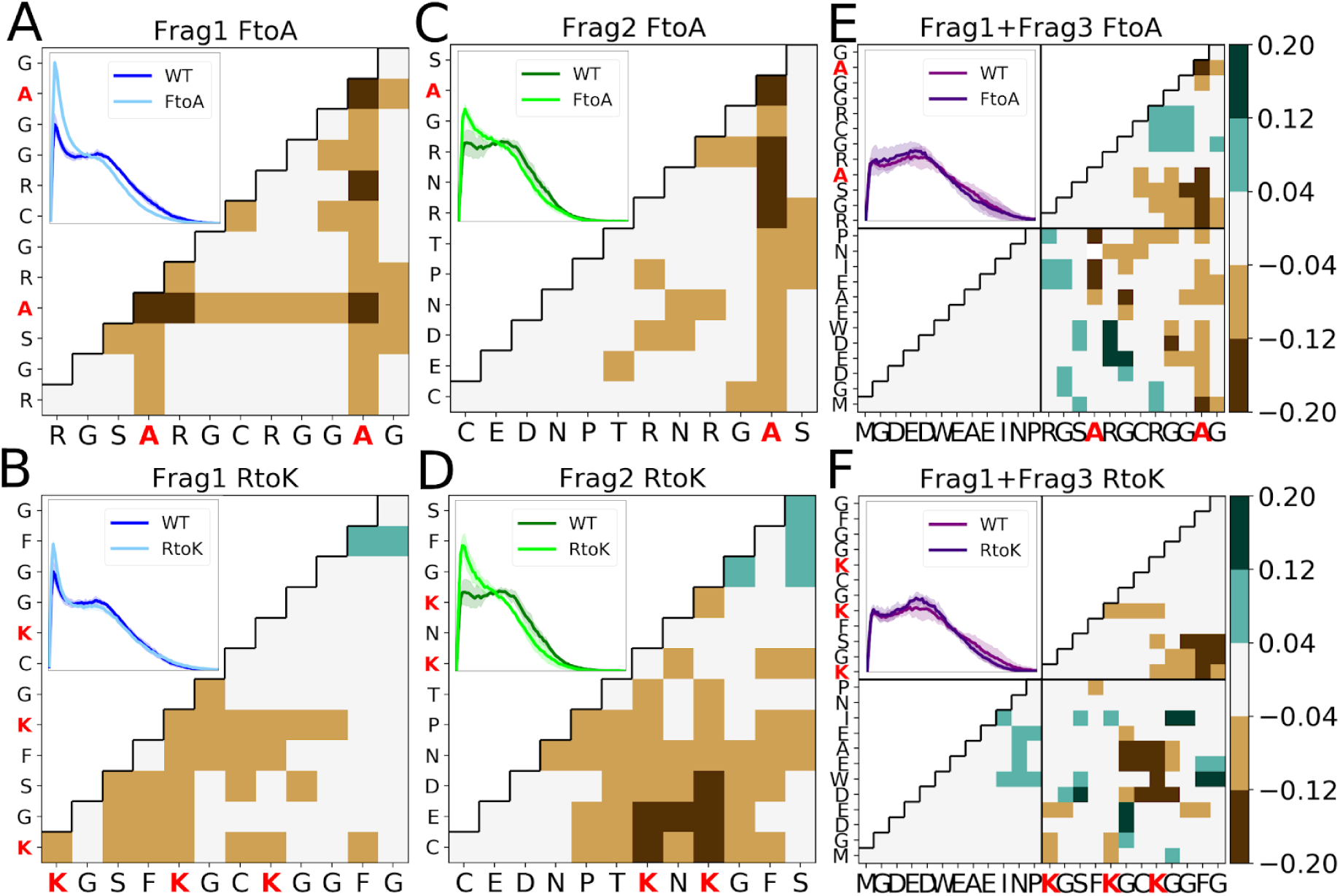
**(A-F)** Variation of the average intermolecular contact probabilities of FtoA (A,C,E) and RtoK (B,D,F) simulations of Frag1, Frag2, and Frag1+Frag3 with respect to wild type (WT) simulations. Positive and negative values indicate respectively an increase and a decrease of the contact probability upon mutation. Insets show the comparison of the distribution of N_C_(i) between WT and mutant (FtoA or RtoK) of the corresponding systems.

Beyond the specific case of DDX4, phenylalanine, tyrosines and arginines are considered key elements in the molecular grammar of cellular LLPS since they have been shown to regulate the phase behaviour of several MLO forming proteins^17,50–52^. Despite this critical role in cellular condensates, our structural understanding of the so-called cation-π and π-π interactions formed by these moieties is currently limited to the statistical analysis of folded domains^25,53,54^ or to the theoretical investigation of extremely simplified models^55^. In this regard, our multi-fragment simulations provide the opportunity to study these interactions in conditions that are more representative of the dynamical and complex environment of liquid condensates. To this aim we focused on Frag1, which contains three arginines and two phenylalanines, and we extended our simulation for a total aggregated time of 10 μs. We first investigated the binding between phenylalanine side chains by determining the free-energy surface as a function of the distance between the centroids of the phenyl groups (D) and the angle ***θ*** between the corresponding planes (Fig. 5A). The resulting landscape is rather shallow and exhibits multiple free-energy minima with similar depth corresponding either to stacked (***θ*** ∼ 0) and T-shaped (60<***θ***<90) conformations. This binding heterogeneity has to be ascribed to the dynamical and crowded environment observed in our simulations. Indeed, the simulation of two isolated phenylalanines resulted into a free-energy surface biased toward stacked conformations (Fig. S5), consistently with the statistical distribution in the protein structure database^53,54^. We then extended this binding mode analysis to phenylalanine-arginine (Fig. 5B) and arginine-arginine (Fig. S6) pairs by monitoring the distance and the angle between guanidinium-phenyl and guanidinium-guanidinium groups, respectively. In both cases, stacked conformations were found to be largely dominant in agreement with the well-established conformational propensities of arginine in protein structures. Interestingly, both Phe-Phe and Arg-Arg free-energy surfaces exhibit less-pronounced minima at intermediate distances (D∼0.75 nm) that cannot be rationalized with direct contact between the side chains and it is hence suggestive of higher-order structuring. We thus analysed our trajectories to probe the formation of transient, intermolecular clusters composed of multiple, interacting phenyl and guanidinium groups. We first characterized the population of these assemblies by reporting in Fig. 5C how Arg and Phe are partitioned in clusters of diverse sizes, ranging from dimers to extended structures involving more than six members. Overall, we found that the roughly half of the Phe and Arg side chains form at least an aromatic-aromatic or aromatic-charged contact (Arg∼40%, Phe∼65%) and that a considerable fraction is involved in higher-order interactions (Arg∼20%, Phe∼30%). Stoichiometry analysis indicated that heterotypic clusters composed of both Phe and Arg side chains are statistically favored over pure assemblies (Fig. 5C). The characterization of the three-dimensional arrangements of phenyl and guanidinium groups revealed a rich structural alphabet. Indeed, analysis of the dimers confirmed preferential stacking of Arg-Arg and Arg-Phe pairs and an equilibrium between parallel and T-shaped conformations in Phe-Phe. These propensities result in complex and dynamic interaction patterns in larger assemblies. Without being exhaustive, we limit here the analysis to trimeric clusters, which correspond to a considerable fraction of the overall population (Arg∼10%, Phe∼16%), and we depict some of their most representative arrangements (Fig. 5D). Among them, sandwich-like structures based on multiple parallel stacking (Fig. 5D top) are particularly common and most often display alternate Arg-Phe-Arg or Phe-Arg-Phe patterns. Furthermore, hybrid trimeric structures combining both T-shaped and parallel interactions (Fig. 5D middle) are frequently observed when two aromatic rings are present whereas triangular architectures (Fig. 5D bottom) are possible albeit less frequent. Structural motifs based on alternate patterns of arginine and aromatic residues can greatly contribute to the thermodynamic stability of protein folds^56,57^. Furthermore, their statistical analysis indicated that even if cation-π-cation interactions are not particularly frequent in protein databases they are widespread and conserved by evolution^56^. Our simulations indicate that intermolecular dynamical interactions with similar and more complex patterns frequently occur between flexible proteins at high concentration. We hypothesize that these higher-order interactions may play an important role in driving the formation of MLOs by promoting the condensation of the scaffolding IDPs/IDRs, which are statistically enriched in arginine and aromatic residues.

**Fig.5.**
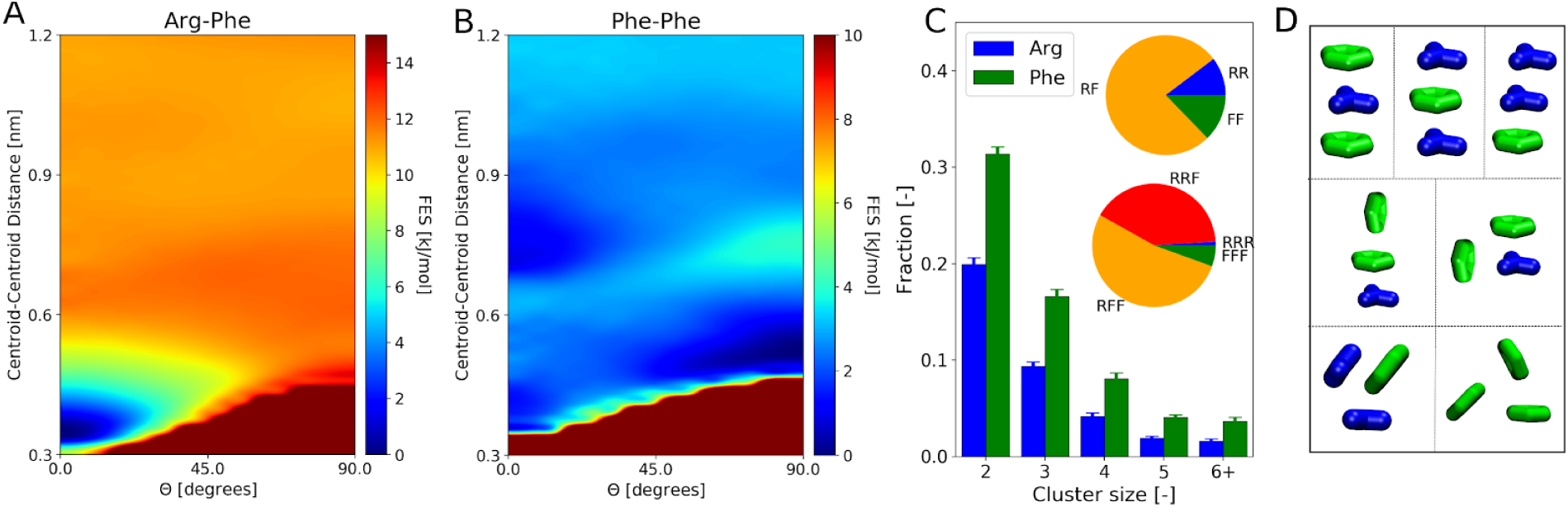
**(A-B)** Free energy surfaces for Phe-Phe (A) and Arg-Phe (B) interactions in Frag 1 simulationas a function of the distances between the centroids of phenyl and guanidinium groups, and their relative orientation theta. **(C)** Histogram of the fraction of either arginines or phenylalanines involved in clusters as a function of the size of the cluster; error bars represent standard deviation across the replicas of Frag1 simulations. In the inset, the average composition of dimers (top) and trimers (bottom). **(D)** Idealized conformations of trimers formed by arginines and phenylalanines, representative of those observed in the MD simulations of Frag1, with multiple stacked conformations (top), combinations of stacked and T-shaped interactions (middle), and triangular patterns (bottom).

The development of accurate models and the increased availability of computational resources have made today atomistic MD simulations a standard and powerful tool in structural biology and molecular biophysics. Nevertheless, the application of this popular approach to the investigation of biomolecular phase separation has lagged behind even if this subject is attracting huge attention in an extremely broad community. In this study, we showed that atomistic MD can be successfully used to unveil the molecular interactions underlying the behaviour of NDDX4 protein, which represents one of the best-characterized models for cellular LLPS. The agreement between our single-molecule simulations of this large-sized IDP and available experimental results reassures us about the accuracy of last-generation force fields and poses the basis for a reliable investigation of NDDX4 intermolecular interactions via a divide-and-conquer strategy. Thus, we focused on the simulation of various protein fragments at high concentration for deciphering how the amino acid sequence modulates their phase behaviour. This approach could be easily extended to perform computational mutagenesis that correctly reproduced the experimental trends and clarified their mechanistic bases. Our results corroborate, with accurate atomistic calculations, previous hypotheses on the correlation between intra- and inter-molecular interactions and on the complex interplay between diverse molecular mechanisms in phase separating systems that were based on lower-resolution simulations. Indeed, our strategy resulted in a high-resolution picture of dynamical intermolecular interactions in conditions mimicking protein condensates. This unprecedented characterization provided access to elusive structural details, such as the transient formation of complex motifs involving multiple arginine and aromatic residues, which are likely to play an important role in stabilizing several MLOs. Considering its generality and affordable computational cost, we are confident that the divide-and-conquer approach proposed here may be fruitfully extended to characterize other phase-separating IDPs. Our future efforts will be devoted to pushing its predictive capabilities with appropriate enhanced sampling techniques in order to further approach the daunting molecular complexity of cellular LLPS.

## Methods

### MD Simulations of NDDX4

Initial configurations of the 236-residue-long NDDX4 were generated with TRaDES-2 software^43,44^ using the sequence reported by Brady et. al^14^. Protein molecules were solvated in a rhombic dodecahedron box with a volume of ∼2290 nm^3^, for a total of approximately 300’000 atoms. The charge of the system was neutralized with 0.1 M of NaCl. The amber99sb-disp force field^40^ was employed to model protein and water molecules. Molecular dynamics (MD) simulations were prepared with the following protocol: 1000 steps of energy minimization with the steepest descent method were carried out to remove unfavorable contacts; water was equilibrated by performing 100 ps of MD simulation in the NVT ensemble followed by 100 ps of MD simulation in the NPT ensemble, in both cases restraining the position of the heavy atoms of the protein molecules, with a force constant of 1000 kJ/(mol nm^2^). Finally, 1 μs of unrestrained MD simulation was performed for each replica, for a cumulated simulation time of 10 μs. All MD simulations were carried out using GROMACS 2018.3^58^. Temperature was kept at 300 K by means of the v-rescale algorithm^59^, controlling separately protein and non-protein molecules in order to avoid the hot-solvent/cold-solute effect^60^. Pressure was kept at 1 bar using the Parrinello-Rahman algorithm^61^. Periodic boundary conditions were applied and long range electrostatic interactions were evaluated using the Particle Mesh Ewald algorithm^62^ with a cutoff of 1 nm for the real space interactions, while van der Waals interactions were computed using a cut off distance of 1 nm. Hydrogen atoms were substituted by virtual sites and all bond lengths were constrained using LINCS^63^, allowing us to use a time step of 4 fs for the integration of the equations of motion^64^. Configurations of the protein were saved every 10 ps and the first 100 ns of each replica were discarded from the analysis. The radius of gyration (R_gyr_) of NDDX4 was computed with the tool gmx gyrate, while the hydrodynamic radius (R_hyd_) was evaluated by means of the HullRad software^65^. Intramolecular contact probabilities were evaluated considering a cut-off of 0.5 nm on the minimum distance between the heavy atoms of pairs of residues (Fig. S7), excluding amino acids closer than four peptide bonds from the calculation. P_loc_(i) variable for each residue *i* was computed by locally averaging the intramolecular contact probabilities of the residues using a rolling window of 7 residues in the interval [i-3,i+3]. Definition of the charged blocks of amino acids and of regions rich in Arg and Phe was performed by analyzing the sequence with a moving window of 11 residues in the interval [i-5,i+5] (Fig. S8). The DSSP algorithm^66^ was employed to estimate the secondary structure propensities from MD simulations, via the interface provided by the gmx do_dssp tool.

### MD Simulations of NDDX4 fragments

Extended structures of the 12-residue-long peptides were generated using the LEaP program included in AmberTools16^67^. Termini of the peptides were capped using ACE and NME groups in order to avoid artificial effects introduced by the charges of the terminal residues. Initial configurations for each replica were generated by inserting the peptides in random positions and orientations in a rhombic dodecahedron box with a volume of ∼145 nm^3^ using the gmx insert-molecules tool available in GROMACS 2018.3^58^. For the Frag1, Frag2, and Frag3 systems, 10 copies of the respectives peptides were inserted. In the case of the Frag1+Frag3 system, 5 copies of each peptide were used. Unlike the simulations of NDDX4, hydrogen atoms were not replaced with virtual sites, considering all the angular degrees of freedom. All bond lengths have been constrained using LINCS allowing a time step of 2 fs. Simulations were then prepared following the same protocol described for the simulations of NDDX4. For each system, three independent replicas were simulated from different initial configurations for 1 μs, except for the WT Frag1 system, where five independent replicas of 2 μs each were simulated. Conformations of the protein molecules have been collected every 10 ps for further analysis. One frame every 1 ns after the first 200 ns was used for the calculation of the intermolecular contact probabilities.

### Structural heterogeneity of NDDX4 fragments complexes

We introduced the variable N_C_(i,t) in order to assess the fluctuations in the local density explored by MD simulations of the fragments selected from NDDX4. This variable was evaluated by means of a switching function with the following form:

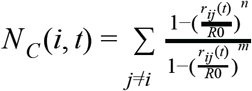

Where r_ij_(t) is the intermolecular distance between alpha carbons at time t, R0 = 1 nm, n=6, and m=12. The evaluation of N_C_(i,t) was performed with an in-house Python3 script which used the MDAnalysis library^68^ to efficiently evaluate the intermolecular distances between alpha carbons. For each system and replica, the P(N_C_) distribution was then computed using all N_C_(i,t) values. One frame every 1 ns was used for the analysis, discarding the first 200 ns of each replica. The P(N_C_) distribution at the density of the condensed phase of NDDX4 (∼380 mg/ml^14^) was computed using 1000 configurations, obtained by placing configurations of Frag1, randomly extracted from the polypeptide MD simulations, at the target concentration by means of the gmx insert-molecules tool.

### Analysis Phe/Arg side chains interactions

Centroid-centroid and minimum distances between heavy atoms of phenyl and guanidinium groups were calculated with PLUMED 2.4.3^69^, while their relative orientations ***θ*** were computed using the gmx gangle tool. The nitrogen atoms of the side chain of arginine and the CD1, CD2, and CZ atoms of phenylalanine were used to define the planes of guanidinium and phenyl groups, respectively. In the evaluation of the free energy surfaces, we chose not to distinguish between angles of ***θ*** and π-***θ***^54^, moreover distance-angle correlation functions were normalised by a factor r^2^sin(***θ***), where r is the centroid-centroid distance, in order to obtain a flat free energy profile in the case of configurations without neither angular nor distance preference. To evaluate the free energy surface relative to the interaction between isolated phenylalanines, we ran four 1-μs-long MD simulations of two uncapped phenylalanines, in a dodecahedron box of volume ∼11 nm^3^. Since the amber99sb-disp does not include zwitterionic residues, we created a new residue type, using the force field parameters of NPHE and CPHE residue types for the ammonium and the carboxyl groups, respectively, while keeping the parameters from the PHE residue type for the rest of the atoms. Charges were then rescaled to have a neutral net charge. We ran the simulations following the same protocol used for the simulations of the polypeptides. In the analysis of higher order structures formed by phenylalanine and arginine, two residues were considered part of the same cluster when at least one pair of heavy atoms belonging to phenyl and guanidinium groups were below the cutoff length of 0.5 nm. The size and the elements belonging to each cluster were evaluated by means of a breadth-first search algorithm implemented in a in-house Python3 script. Configurations every 100 ps of the Frag1 simulations were used for the analysis and structures of the trimers were finally extracted using the gmx trjconv tool.

## Supplementary Information

**Fig.S1.**
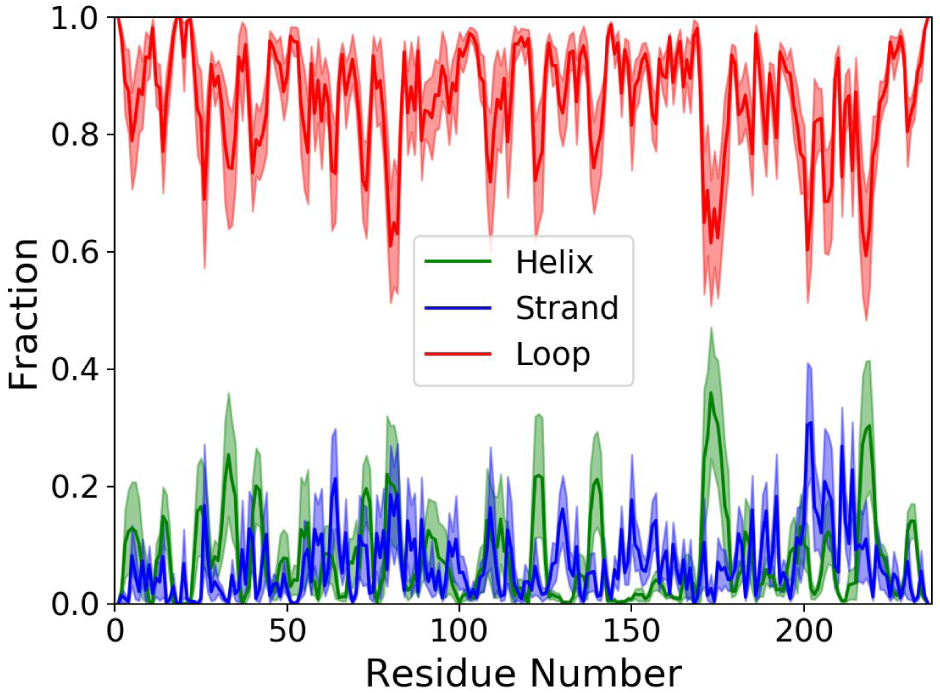
Average secondary structure propensity from MD simulations of NDDX4, shaded areas indicate the error of the mean over 10 replicas.

**Fig.S2.**
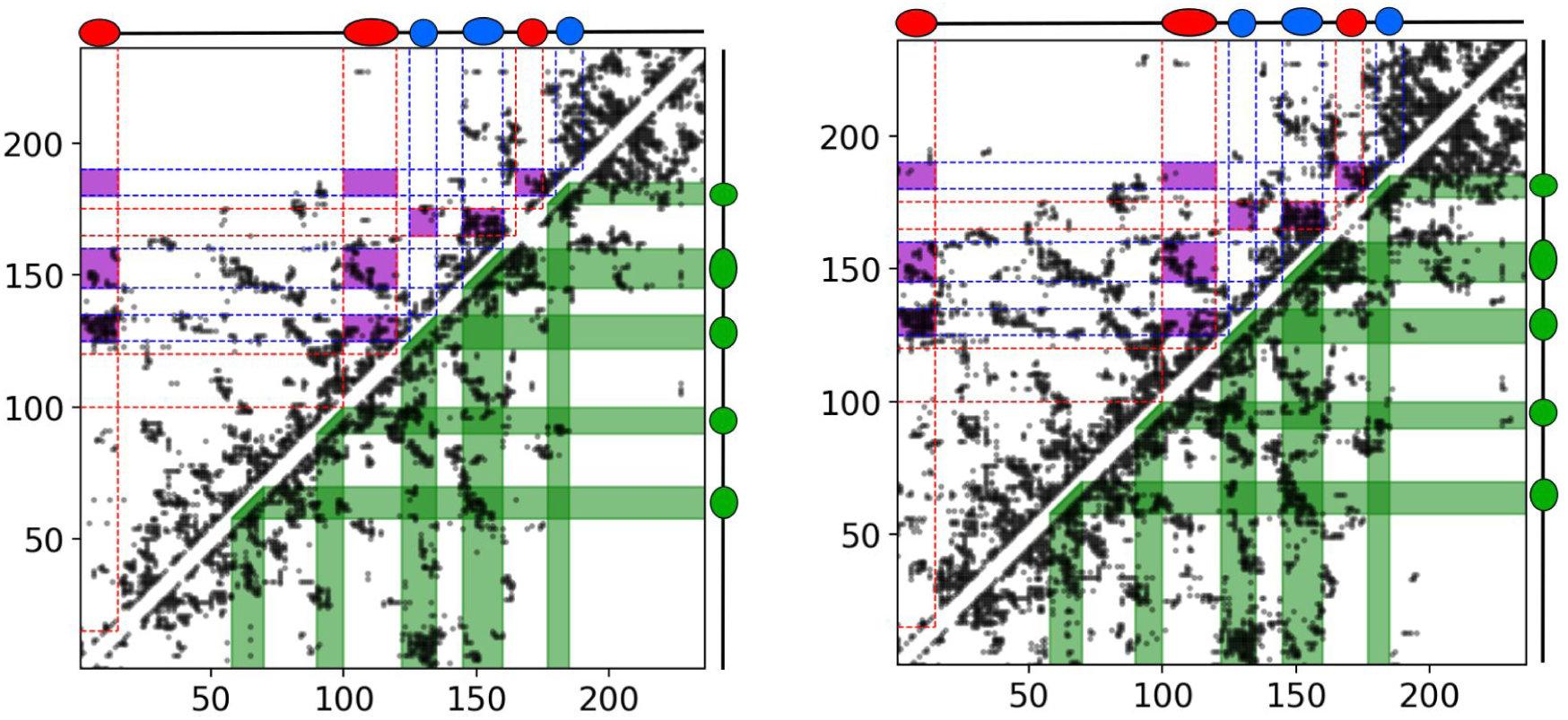
Residue-residue contact probability map as in Fig.1D. Scatter plots represent, respectively, the 10% (left) and 15% (right) most frequent contacts observed in the MD simulations.

**Fig.S3.**
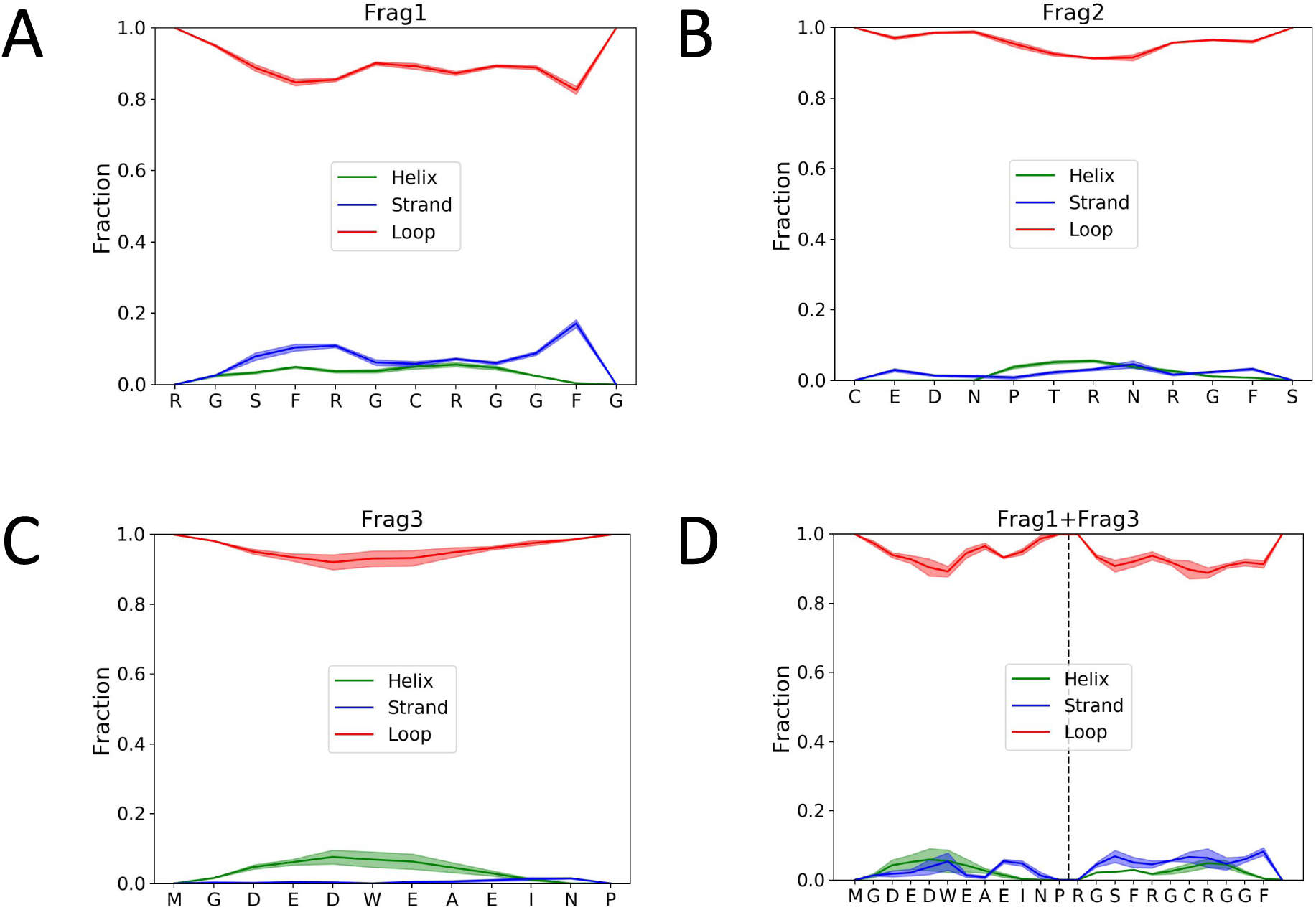
**(A-D)** Average secondary structure propensity from MD simulations of the simulated fragments. Shaded areas indicate the error of the mean over the replicas.

**Fig.S4.**
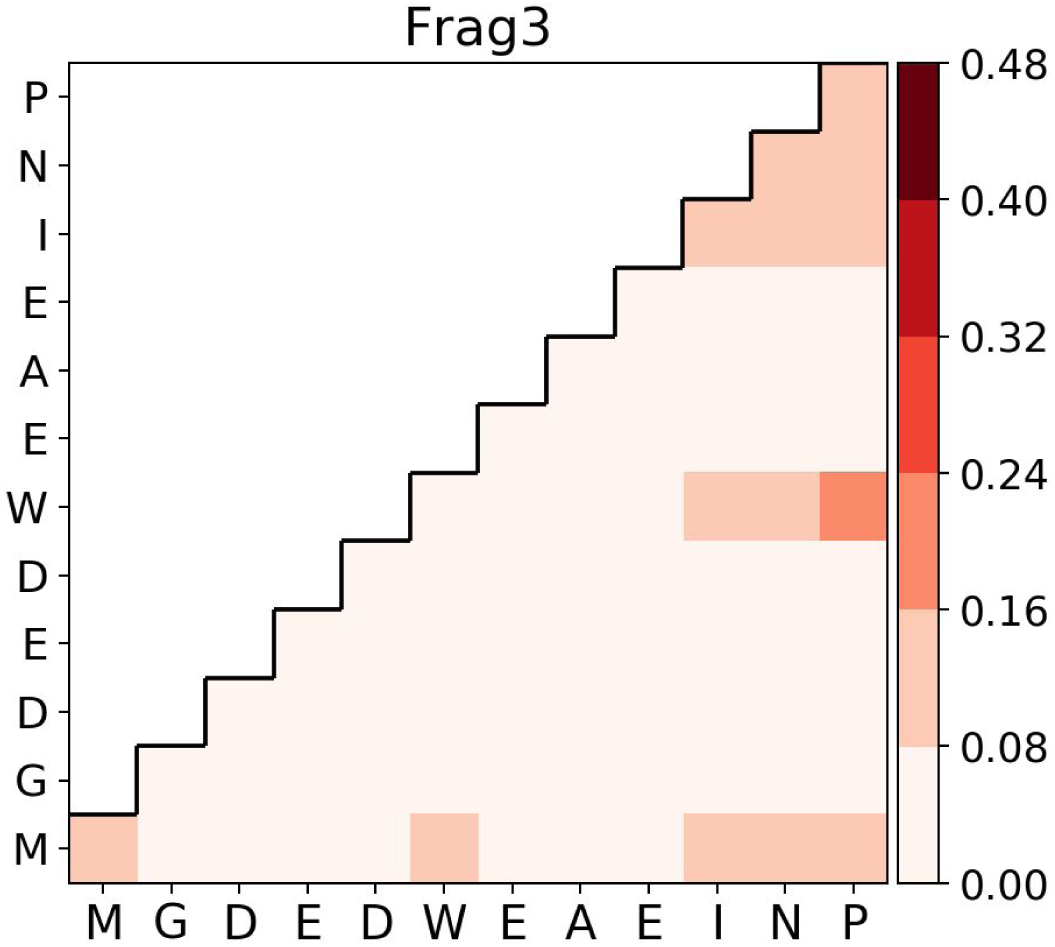
Average intermolecular contact probability from Frag3 simulations.

**Fig.S5.**
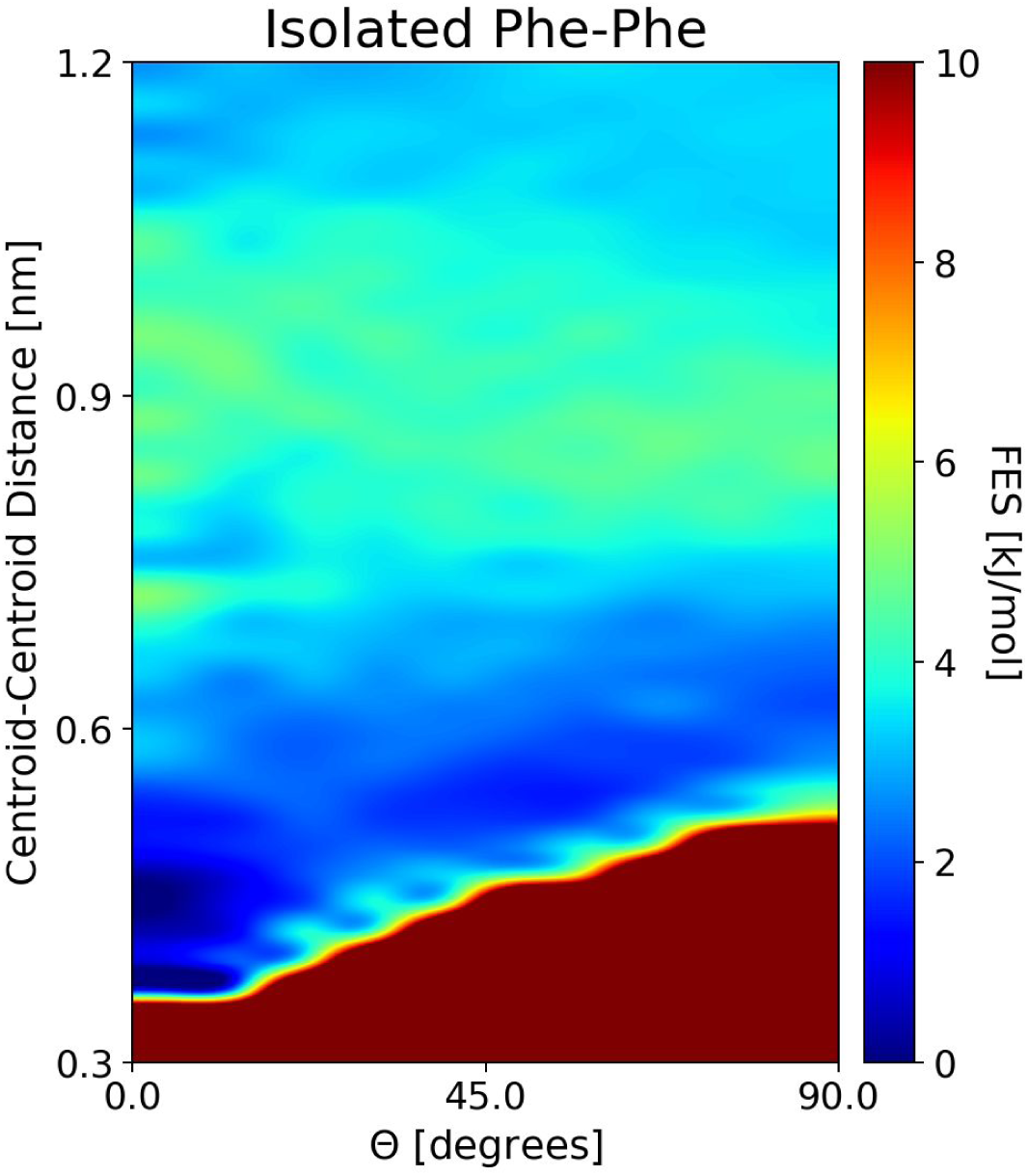
Free energy surfaces for isolated Phe-Phe interactions as a function of the distances between the centroids of guanidinium groups, and their relative orientation theta.

**Fig.S6.**
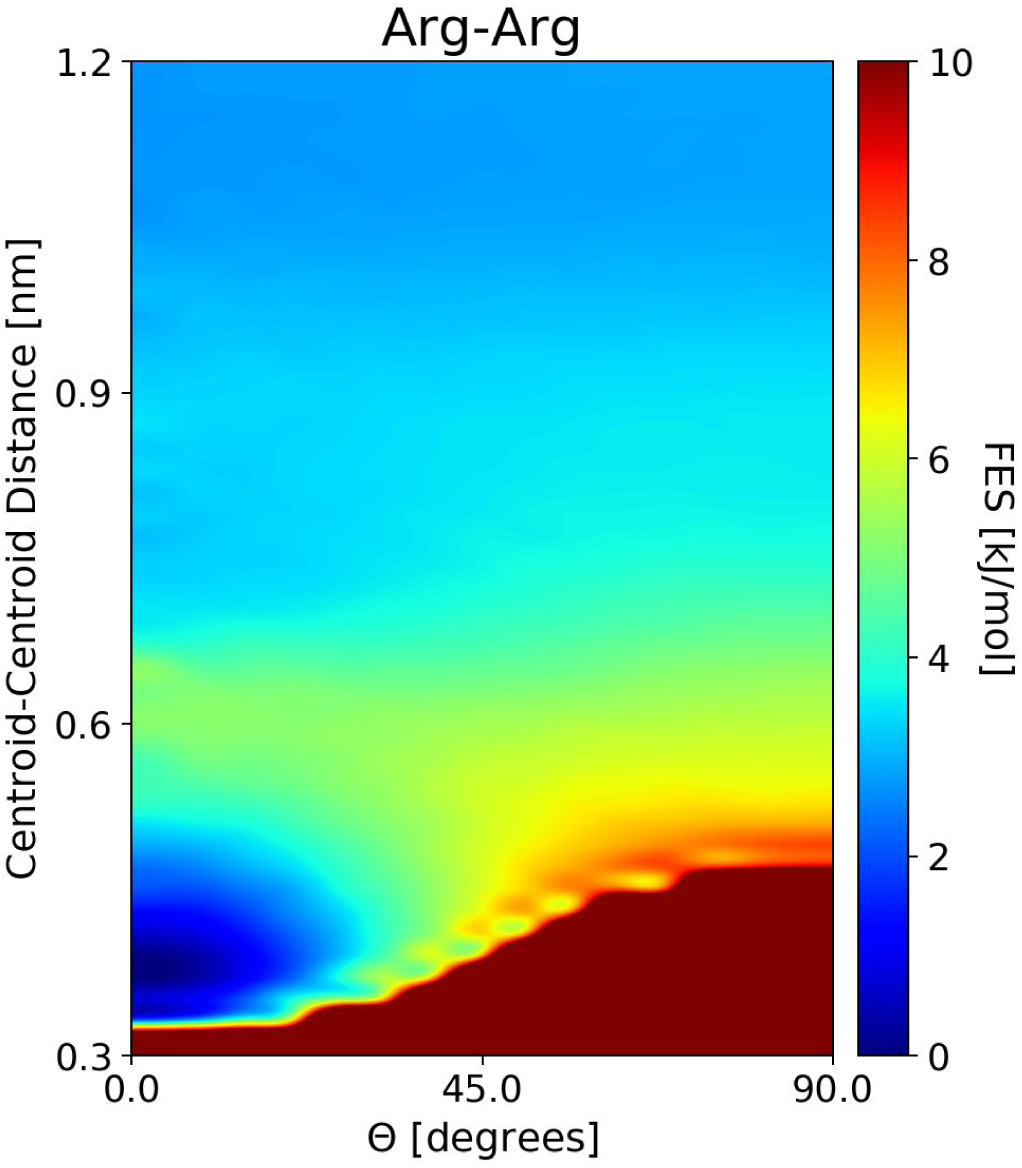
Free energy surfaces for Arg-Arg interactions as a function of the distances between the centroids of guanidinium groups, and their relative orientation theta.

**Fig.S7.**
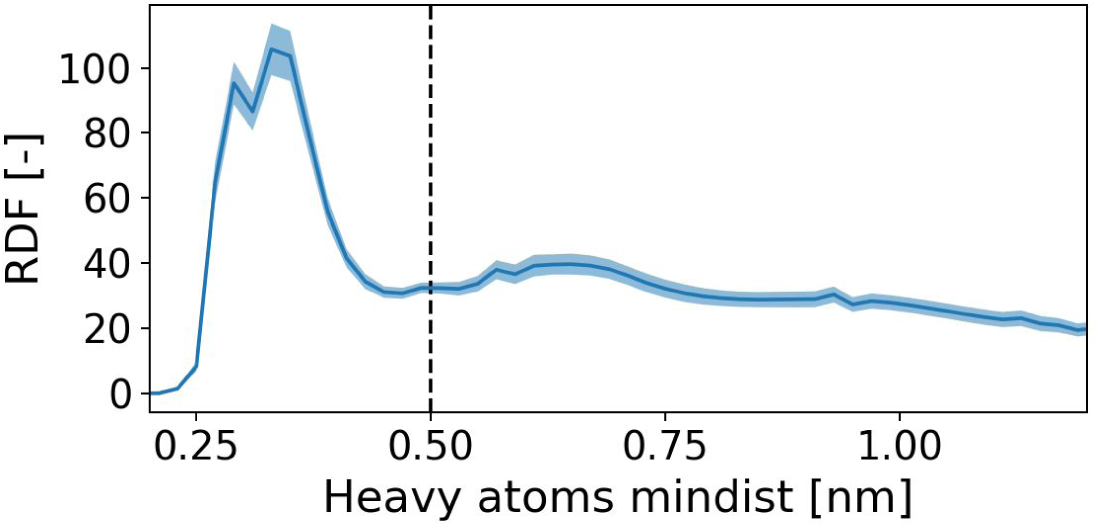
Average inter-residue radial distribution function (RDF) of the minimum distance between heavy atoms from MD simulations of NDDX4. Shaded area represents standard deviation over the replicas. The cut-off used for the definition of the contacts (0.5 nm) is indicated by the dashed line.

**Fig.S8.**
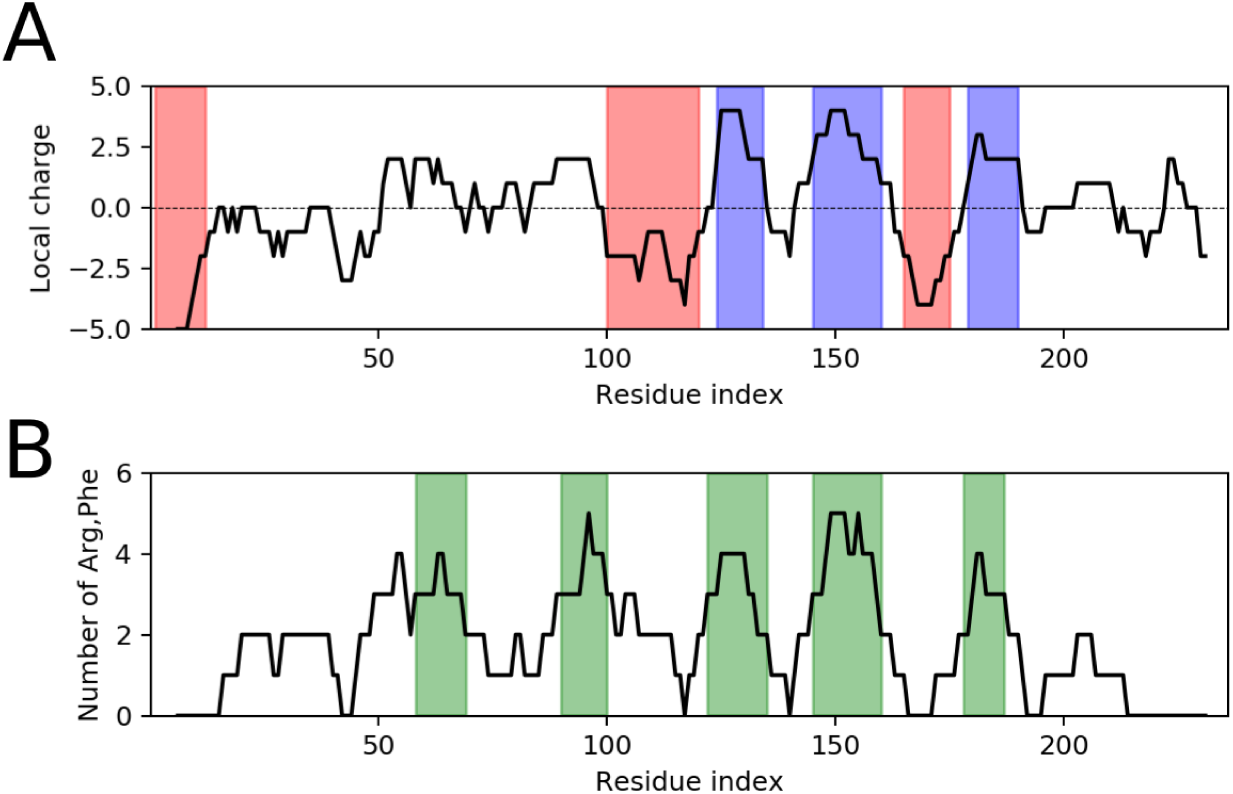
**(A)** Local charge along the NDDX4 sequence using a moving window of 11 residues. Blue and red shaded areas represent respectively the position of the positively and negatively charged blocks used in Fig.1 and Fig.S3. **(B)** Number of arginine and phenylalanine residues along the sequence of NDDX4 using a moving window of 11 residues. Green shaded areas represent the blocks used in Fig.1D and Fig.S3.

